# Clustering and compositionality of task representations in a neural network trained to perform many cognitive tasks

**DOI:** 10.1101/183632

**Authors:** Guangyu Robert Yang, H. Francis Song, William T. Newsome, Xiao-Jing Wang

## Abstract

A neural system has the ability to flexibly perform many tasks, but the underlying mechanism cannot be elucidated in traditional experimental and modeling studies designed for one task at a time. Here, we trained a single network model to perform 20 cognitive tasks that may involve working memory, decision-making, categorization and inhibitory control. We found that after training, recurrent units developed into clusters that are functionally specialized for various cognitive processes. We introduce a measure to quantify relationships between single-unit neural representations of tasks, and report five distinct types of such relationships that can be tested experimentally. Surprisingly, our network developed compositionality of task representations, a critical feature for cognitive flexibility, whereby one task can be performed by recombining instructions for other tasks. Finally, we demonstrate how the network could learn multiple tasks sequentially. This work provides a computational platform to investigate neural representations of many cognitive tasks.

## INTRODUCTION

The prefrontal cortex is important for numerous cognitive functions^1–3^, partly because of its central role in task representation^4–7^. Electrophysiological experiments using behaving animals reported prefrontal neurons that are either selective for different aspects of a given task^8,9^ or functionally mixed^10,11^. Much less is known about functional specialization of task representations at the neuronal level. Imagine a single-neuron recording that could be carried out with animals switching between many different tasks. Is each task supported by a “private” set of neurons, or does each task involve every neuron in the network, or somewhere in between? If two tasks require a common underlying cognitive process, such as working memory or decision making, what would be the relationship between their neural representations? In other words, what would be the “neural relationship” between this pair of tasks? Would the two tasks utilize a shared neural substrate?

Humans readily learn to perform many cognitive tasks in a short time. By following verbal instructions such as “Release the lever only if the second item is not the same as the first,” humans can perform a novel task without any training at all^6^. A cognitive task is typically composed of elementary sensory, cognitive, and motor processes^5^. Performing a task without training requires composing elementary processes that are already learned into temporal sequences that enable correct performance on the new task. This property, called “compositionality,” has been proposed as a fundamental principle underlying flexible cognitive control^12^. Indeed, human studies have suggested that the representation of complex cognitive tasks in the lateral prefrontal cortex is compositional^16,13^. However, these tasks involved verbal instructions; it is unknown whether non-verbal tasks commonly used in animal physiological experiments also display compositionality and whether relatively simple neural network models are sufficient to support compositional task structures.

These questions remain difficult to address with conventional experimental and modeling approaches. Experiments with laboratory animals have so far been largely limited to a single task at a time; on the other hand, human imaging studies lack the spatial resolution to address questions at the single neuron level. Therefore, the lack of neural recordings from animals performing many different tasks leaves unanswered important questions regarding how a single network represents and supports distinct tasks. In principle, these questions could be addressed in neural circuit models, but designing a single neural circuit model capable of multiple tasks is challenging and virtually nonexistent. To tackle these problems, we took the approach of training recurrent neural networks (RNNs)^11,14–19^. In this work, we trained a single RNN to perform 20 cognitive tasks. We found that after training, the emerging task representations are organized in the form of clustering of recurrent units. Our network also makes numerous testable predictions regarding the relationship between the neural representations of pairs of cognitive tasks. Surprisingly, we found that compositionality of task representations emerges from training in our network model, which can be instructed to perform new tasks without further training. Our work provides a framework for investigating neural representations of task structures and neural relationships between tasks.

## RESULTS

### Training neural networks for many cognitive tasks

To study how various cognitive tasks might be implemented in a single neural circuit, we trained a recurrent neural network model (Fig. 1a) to perform 20 tasks, most of which are commonly used in neurophysiological studies of nonhuman animals and crucial to our understanding of the neural mechanisms of cognition. The chosen set of tasks includes variants of memory-guided response^20^, simple perceptual decision making^21^, context-dependent decision-making^11,22^, multi-sensory integration^23^, parametric working memory^24^, inhibitory control (e.g., in anti-saccade)^25^, delayed match-to-sample^26^, and delayed match-to-category^27^ tasks (Table 1, Supplementary Fig. 1).

The recurrent network model emulates a “cognitive-type” cortical circuit such as the pre-frontal cortex^3^, which receives converging inputs from multiple sensory pathways and projects to downstream motor areas. We designed our network architecture to be general enough for all the tasks mentioned above, but otherwise as simple as possible to facilitate analysis. For every task, the network receives noisy inputs of three types: fixation, stimulus, and rule (Fig. 1a). The fixation input indicates whether the network should “fixate” or respond (e.g. “saccade”). Thus the decrease in the fixation input provides a “go signal” to the network. The stimulus inputs consist of two modalities, each represented by a ring of input units that encodes a one-dimensional circular variable such as motion direction or color on a color wheel^18^. A single rule input unit is activated in each trial, instructing the network on which task it is currently supposed to perform. The network projects to a fixation output unit and a group of motor units encoding the response direction as a one dimensional variable on a ring of outputs (e.g., saccade direction, reach direction). To mimic biological neurons, all units in our recurrent network receive private noise and have non-negative activities, imposed by a realistic neuronal input-output function^28^.

Before training, a network is incapable of performing any task. It is trained with supervised learning^11,15^, which modifies all connection weights (input, recurrent, and output) to minimize the difference between the network output and a desired (target) output. All tasks were randomly interleaved during training (at the end we will present results from sequential training). Below we show results obtained from networks of 256 recurrent units, and results are robust with re-spect to the exact network size. After training, a single network model achieved high behavioral performance across all tasks (Fig. 1b). Furthermore, by conducting a battery of psychometric tests, we demonstrate that the network displays behavioral features consistent with animal studies. For instance, in perceptual decision-making tasks, the network achieves better performance with higher coherence and longer duration of the stimulus (Fig. 1c, Supplementary Fig. 2a-f)^21^, and it combines information from different sources to form decisions (Fig. 1d)^23^. In working memory tasks, the network can maintain information throughout a delay period of up to five seconds (Supplementary Fig. 2g)^1,20,24^.

**Figure 1.**
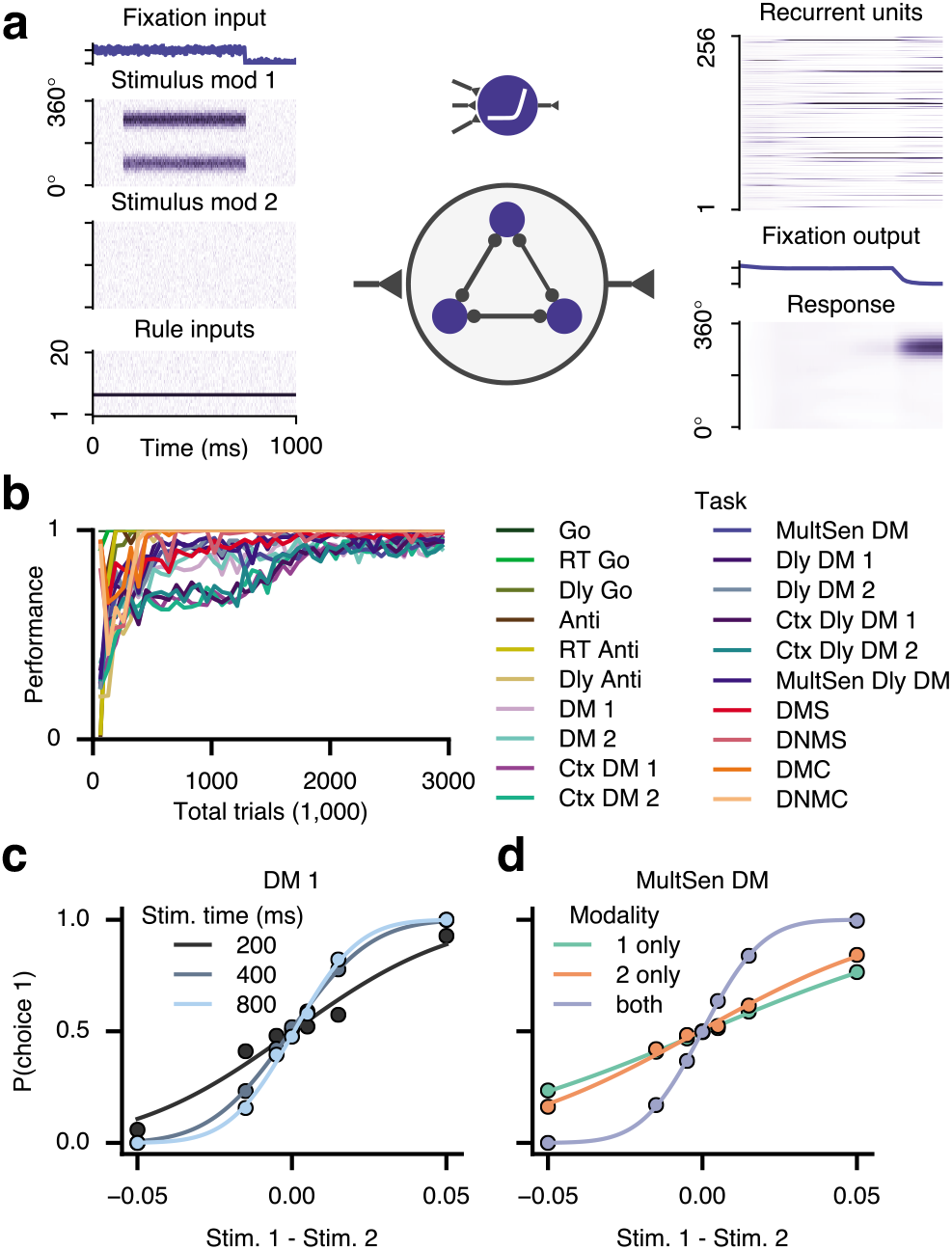
A recurrent neural network model is trained to perform a large number of cognitive tasks. **(a)** A recurrent neural network (middle) described by rate units receives inputs (left) encoding a fixation cue, stimuli from two modalities, and a rule signal (which instructs the system which task to perform in a given trial). The network has 256 recurrent units (top right), and it projects to a fixation output unit (which should be active when a motor response is unwarranted) and a population of units selective for response directions (right). All units in the recurrent network have non-negative firing rates. All connection weights and biases are modifiable by training using a supervised learning protocol. **(b)** The network successfully learned to perform 20 tasks. **(c, d)** Psychometric curves in two decision making (DM) tasks. **(c)** Perceptual decision-making relies on temporal integration of information, as the network performance improves when the noisy stimulus is presented for a longer time. **(d)** In a multi-sensory integration task, the trained network combines information from two modalities to improve performance (compared with performance when information is only provided by a single modality).

**Table 1.** Names and abbreviations of all tasks trained in the networks. Most of the trained tasks are derived from archetypal cognitive tasks used in non-human animal experiments. We grouped our tasks into five task families. We are not aware of experimental studies that investigated the Ctx Dly DM 1, Ctx Dly DM 2, or MultSen Dly DM tasks in non-human animals.

| Task name | Abbreviation | Task family | Reference |
| --- | --- | --- | --- |
| Go | Go | Go | N/A |
| Reaction-time go | RT Go | Go | N/A |
| Delayed go | Dly Go | Go | [20] |
| Anti-response | Anti | Anti | [25] |
| Reaction-time anti-response | RT Anti | Anti | [25] |
| Delayed anti-response | Dly Anti | Anti | [25] |
| Decision making 1 | DM 1 | DM | [21] |
| Decision making 2 | DM 2 | DM | [21] |
| Context-dependent decision making 1 | Ctx DM 1 | DM | [11] |
| Context-dependent decision making 2 | Ctx DM 2 | DM | [11] |
| Multi-sensory decision making | MultSen DM | DM | [23] |
| Delayed decision making 1 | Dly DM 1 | Dly DM | [24] |
| Delayed decision making 2 | Dly DM 2 | Dly DM | [24] |
| Context-dependent delayed decision making 1 | Ctx Dly DM 1 | Dly DM | N/A |
| Context-dependent delayed decision making 2 | Ctx Dly DM 2 | Dly DM | N/A |
| Multi-sensory delayed decision making | MultSen Dly DM | Dly DM | N/A |
| Delayed match-to-sample | DMS | Matching | [26] |
| Delayed non-match-to-sample | DNMS | Matching | [26] |
| Delayed match-to-category | DMC | Matching | [27] |
| Delayed non-match-to-category | DNMC | Matching | [27] |

### Dissecting the circuit for the family of Anti tasks

For trained neural networks to be useful model systems for neuroscience, it is critical that we attempt to understand the circuit mechanism underlying the network computation^29^. Here we demonstrate how a trained network could be dissected and analyzed in a sample family of cognitive tasks. Anti-response tasks are important tools to investigate voluntary action and inhibitory control^25^. These tasks require an anti-response, in the opposite direction from the more common pro-response towards a stimulus’ location. Our set of tasks includes three tasks from the Anti task family (Table 1): the anti-response (Anti), reaction-time anti-response (RT Anti), and delayed anti-response (Dly Anti) tasks. We found that a subgroup of units emerged in a trained network, which we call Anti units (Fig. 2a). These units are primarily selective to stimuli in the Anti family of tasks. Inactivating or “lesioning” all Anti units at once resulted in a complete failure in performing the family of tasks that require an anti-response, but had essentially no impact on the performance of the other tasks (Fig. 2b).

Since we have access to all the information of the trained network, we next investigated the connection weights of Anti units to understand their roles. Anti units receive positive connection weights from the three rule input units representing Anti tasks (Fig. 2c), which explained why Anti units are only active during Anti tasks. Next, we studied the connection weights of Anti units with the stimulus-encoding input ring and the response-encoding output ring. For each Anti unit, the preferred input and output directions defined by the input and output connection weights are 180 degrees apart (Fig. 2d). These opposite preferred directions serve as the neural substrate for vector inversion (anti-mapping) required by Anti tasks. Finally, the Anti units strongly inhibit the rest of the recurrent units (Non-Anti units) through recurrent connections (Fig. 2e), suppressing a pro-response with inhibitory control. Thus, the circuit mechanism underlying Anti tasks in our trained network is delineated: A group of units emerge from training that are specialized for the anti-response process and are essential in every task that requires this process. The Anti rule inputs engage vector-inverting Anti units, which in turn exert inhibitory control over Non-Anti units (Fig. 2f).

**Figure 2.**
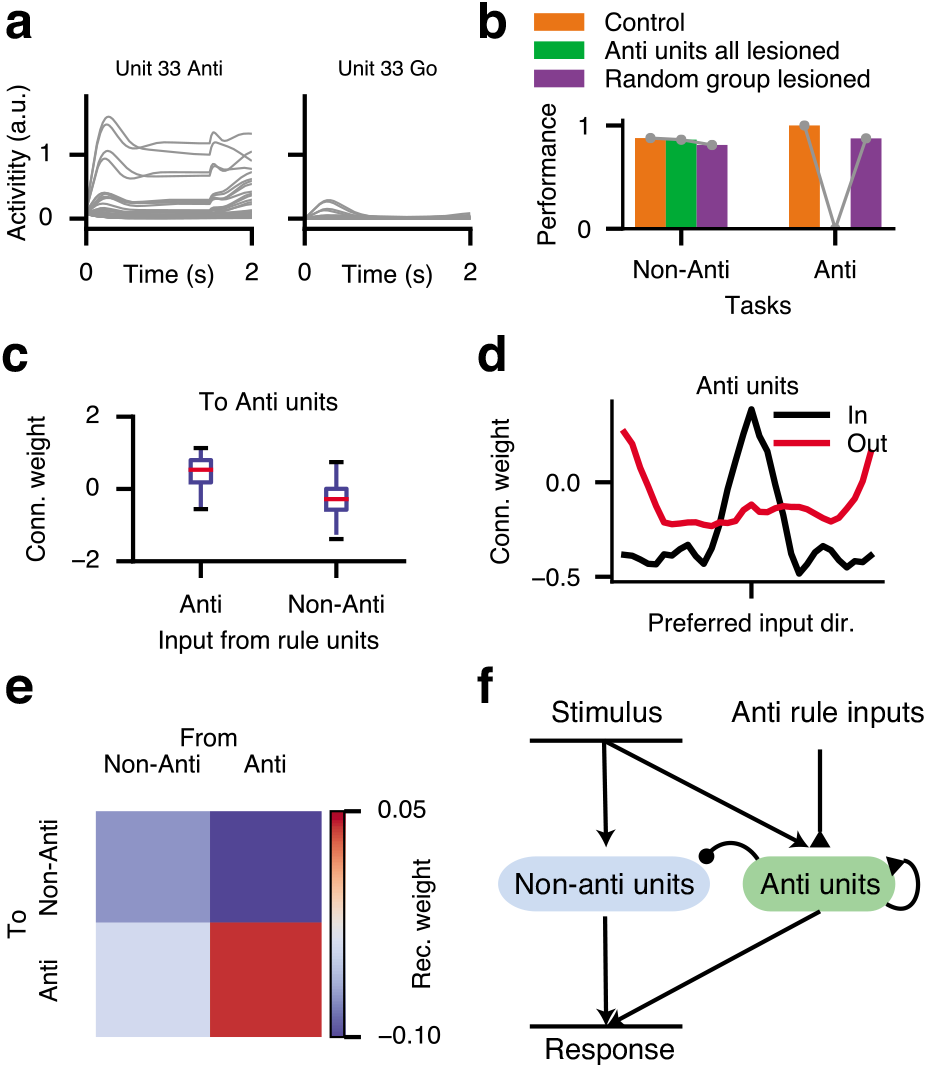
Dissecting the circuit for a family of tasks. **(a)** An example Anti unit, which is primarily selective in the Anti-family of tasks. Different traces show neural activities across stimulus conditions within a task. **(b)** After lesioning all Anti units together (green), the network can no longer perform any of the Anti tasks, while performance for other tasks remain intact. Instead, lesioning the same number of randomly selected units had a minor impact on the performance. **(c)** Anti units receive strong positive connections from rule units representing the Anti tasks but negative connections from non-Anti rule units. The box shows the lower to upper quartiles of connection weights. The whiskers show the full range. **(d)** Average connections from input units to Anti units (black) and those onto output units (red) display opposite preferred directions, thereby vector conversion (from pro- to anti-response) is realized. Both input and output connections are sorted by each unit’s preferred input direction, defined as the stimulus direction represented by the strongest-projecting input unit. **(e)** Network wiring architecture that emerged from training, in which Anti units excite themselves and strongly inhibit other units. **(f)** Circuit diagram summarizing the neural mechanism of the Anti-family tasks.

### Functional clusters encode subsets of tasks

The focus of our analysis was to examine the neural representation of tasks. After training, it is conceivable that each unit of the recurrent network is only selective in one or a few tasks, forming highly-specialized task representations. On the other hand, task representations may be completely mixed, where all units are engaged in every task. We sought to assess where our network lies on the continuum between these two extreme scenarios.

To quantify single-unit task representation, we need a measure of task selectivity that is general enough so it applies to a broad range of tasks, and at the same time simple enough so it can be easily computed. We propose a measure that we call Task Variance (see Online Methods). For each unit, the task variance for a given task is obtained by first computing the variance of neural activities across all possible task conditions at a given time point, then averaging that variance across time (excluding the fixation epoch) (Fig. 3a). Task variance is agnostic about the task setup and can be easily computed in models and is also applicable to the analysis of experimental data.

By computing the task variance for all trained tasks, we can study how individual units are differentially selective in all the tasks (Fig. 3b). For better comparison across units, we normalized the task variance of each unit such that the maximum normalized variance over all tasks is one. By analyzing the patterns of normalized task variance for all active units, we found that units are self-organized into distinct clusters through learning (Fig. 3c,d) (see Online Methods). We identified about 10 clusters in the network. Each cluster is mainly selective in a specific subset of tasks. To understand the causal role of these clusters, we lesioned each of them while monitoring the change in performance across all 20 tasks (Fig. 3e). We found one cluster (cluster number 3) that is specialized for the Anti-family tasks, and it consists mainly of Anti units analyzed in Fig. 2. Another two clusters (cluster numbers 6 and 7) are specialized for decision-making tasks involving modality 1 and 2 respectively. Furthermore, one cluster (cluster number 5) selective in the parametric working memory tasks (Dly DM task family) is also selective in the perceptual decision making tasks (DM task family), indicating a common neural substrate for these two cognitive functions in our network^30^. We can also study how units are clustered based on epoch variance, a measure that quantifies how selective units are in each task epoch (Supplementary Fig. 3). One cluster of units presumably supports response generation, as it is highly selective in the response epoch but not the stimulus epoch. Our results indicate that the network successfully identified common sensory, cognitive, and motor processes underlying subsets of tasks, and through training developed units dedicated to the shared processes rather than the individual tasks.

**Figure 3.**
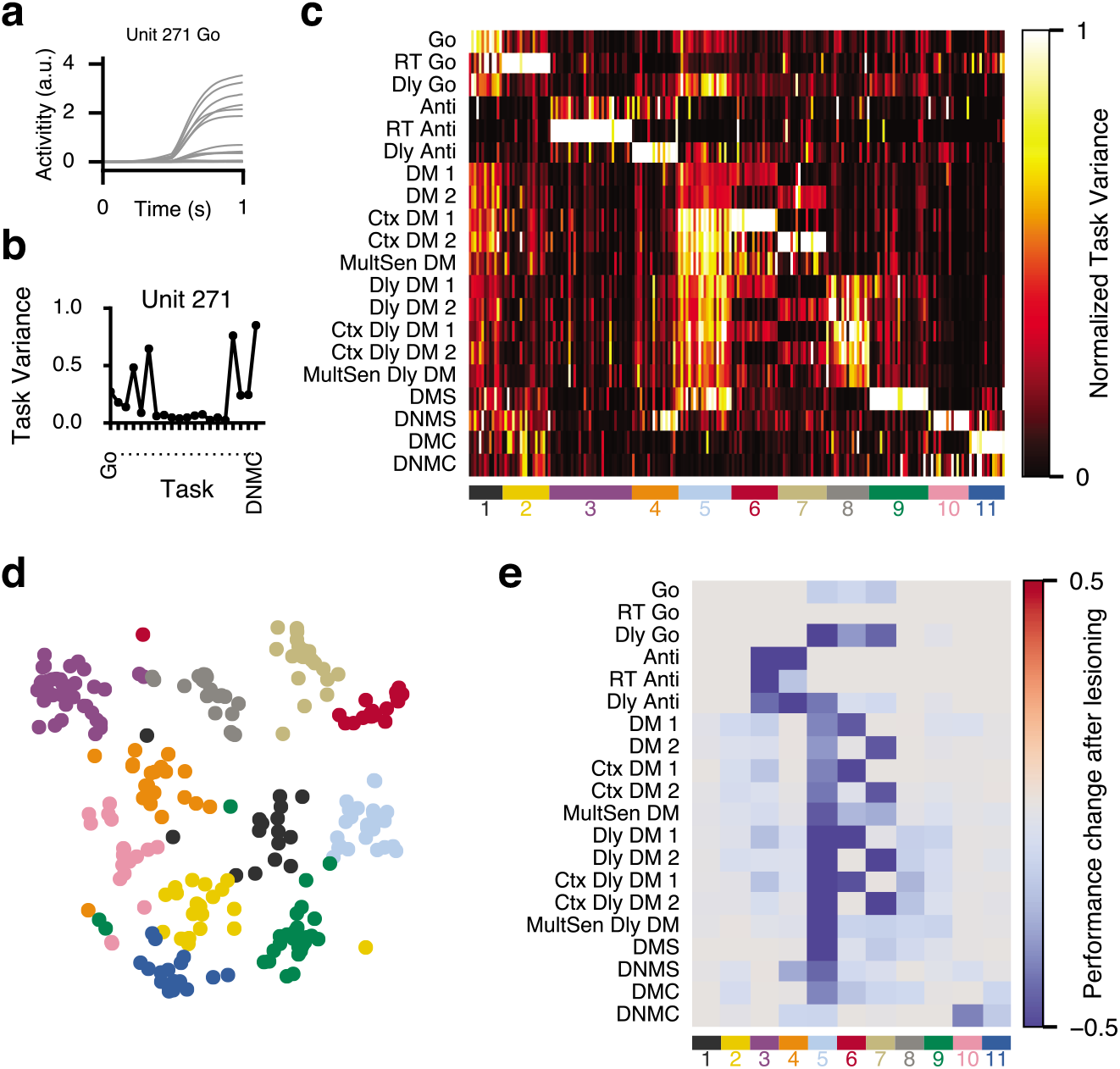
The emergence of functionally specialized clusters for task representation. **(a)** Neural activity of a single unit during an example task. Different traces correspond to different stimulus conditions. **(b)** Task variances across all tasks for the same unit. For each unit, task variance measures the variance of activities across all stimulus conditions. **(c)** Task variances across all tasks and active units, normalized by the peak value across tasks for each unit. Units form distinct clusters identified using the K-means clustering method based on normalized task variances. Each cluster is specialized for a subset of tasks. A task can involve units from several clusters. Units are sorted by their cluster membership, indicated by colored lines at the bottom. **(d)** Visualization of the task variance map. For each unit, task variances across tasks form a vector that is embedded in the two-dimensional space using t-distributed Stochastic Neighbor Embedding (t-SNE). Units are colored according to their cluster membership. **(e)** Change in performance across all tasks when each cluster of units is lesioned.

### Relationships between neural representations of pairs of tasks

The map of normalized task variance in Fig. 3c allowed us to visualize the whole network across many tasks all at once. However, it is of limited use when we try to compare with experimental data or to analyze the (dis)similarity of the neural task representation between any pair of tasks. To quantify how each unit is selective in one task in comparison to another task, we introduce a simple measure based on task variance: the Fractional Task Variance (FTV). For unit *i*, the fractional task variance with respect to task A and task B is defined as

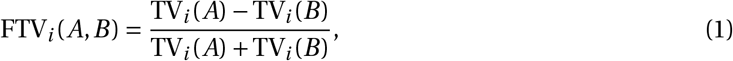

where TV*_i_(A)* and TV*_i_(B)* are the task variances for tasks A and B respectively. Fractional task variance ranges between −1 and +1. Having a FTV*_i_(A,B)* close to +1 (or −1) means that unit *i* is primarily selective in task A (or B).

For every pair of tasks, we can compute the fractional task variance for all units that are active in at least one of the two tasks. Each distribution of FTVs contains rich information about the single-unit level neural relationship between the pair of tasks. Having 20 tasks provides us with 190 distinct FTV distributions (Supplementary Fig. 4), from the shape of which we summarized five typical neural relationships (Fig. 4).

1. Disjoint (Fig. 4a). When two tasks have a disjoint relationship like the Anti task and the DM1 task, the FTV distribution is characterized by two peaks at the two ends and few units in between. There is little overlap between units selective in the two tasks. The shape of the FTV distribution is rather robust across independently trained networks: The FTV distribution from one sample network closely matches the averaged distribution from 20 networks.
2. Inclusive (Fig. 4b). This relationship is embodied by a strongly skewed FTV distribution, suggesting that one task is neurally a subset of another task. In this case, there are no units that are selective in the DM1 task yet not in the Dly DM 1 task.
3. Mixed (Fig. 4c). A mixed relationship is characterized by a broad uni-modal FTV distribution centered around 0 with no clear peak at the two ends. This distribution suggests that the two tasks utilize overlapping neural circuits.
4. Disjoint-Equal (Fig. 4d). For Ctx DM 1 and 2, the FTV distribution is trimodal, with two peaks at the two ends and an additional peak around 0. This relationship can be considered as a combination of the Disjoint relationship and the Equal relationship. The Equal relationship is represented by a single, narrow peak around 0. In this scenario, the two tasks each gets a private neural population, while they also share the third population.
5. Disjoint-Mixed (Fig. 4e). This relationship is a combination of the Disjoint and the Mixed relationships. Many units only participate in one of the two tasks, while the rest of the units are mixed in both tasks.

**Figure 4.**
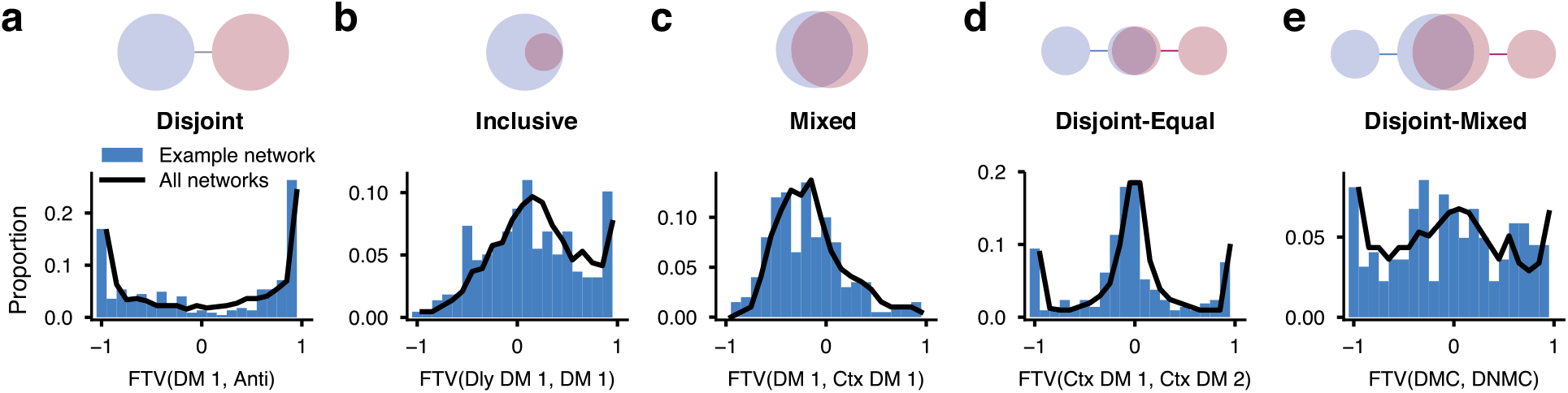
A diversity of neural relationships between pairs of tasks. For a pair of tasks, we characterize their neural relationship by the distribution of fractional task variances over all units. We observed five typical relationships: Disjoint **(a)**, Inclusive **(b)**, Mixed **(c)**, Disjoint-Equal **(d)**, and Disjoint-Mixed **(e)**. Blue: distribution for one example network. Black: averaged distribution over 20 networks.

In summary, we introduced a simple yet informative measure to study the diverse neural relationships between pairs of tasks. We found that these relationships can be categorized into several canonical types. Our results on FTV distributions (Supplementary Fig. 4) provide an array of straightforward predictions on pairwise neural relationships between cognitive tasks.

### Compositional representations of tasks

A cognitive task can, in general, be expressed abstractly as a sequence of sensory, cognitive and motor processes, and cognitive processes may involve a combination of basic functions (such as working memory) required to perform the task. The compositionality of cognitive tasks is natural for human subjects because tasks are instructed with natural languages, which are compositional in nature^12^. For example, the Go task can be instructed as “Saccade to the direction of the stimulus after the fixation cue goes off,” while the Dly Go task can be instructed as “Remember the direction of the stimulus, then saccade to that direction after the fixation cue goes off.” Therefore, the Dly Go task can be expressed as a composition of the Go task with a particular working memory process. Similarly, the Anti task can be combined with the same working memory process to form the Dly Anti task.

Here we test whether the network developed compositional representations for tasks, even when it was never explicitly provided with the relationships between tasks. For the sake of simplicity, we studied the representation of each task as a single high-dimensional vector. To compute this “task vector”, we averaged neural activities across all possible stimulus conditions within each task and focused on the steady-state response during the stimulus epoch (Fig. 5a). Most tasks studied here begin with a stimulus epoch, so the neural population state near the end of stimulus presentation is potentially representative of how the network processed the stimulus in a particular task to meet the computational need of subsequent behavioral epochs. Indeed, this idea is confirmed using principal component analysis, which revealed that task vectors in the state space spanned by the top two principal components are distinct for all twenty tasks (Supplementary Fig. 5).

When plotting the task vectors representing the Go, Dly Go, Anti, and Dly Anti tasks, we found that the vector pointing from the Go vector towards the Dly Go vector is very similar to the vector pointing from the Anti vector to the Dly Anti vector (Fig. 5b). This finding is surprisingly robust and becomes even more apparent when we combined results from many networks (Fig. 5c). The Go-to-Dly Go vector and the Anti-to-Dly Anti vector presumably reflect the cognitive process of working memory. Similar findings are made with another set of tasks. The vector pointing from the Ctx DM 1 task to the Ctx DM 2 task is similar to the vector pointing from the Ctx Dly DM 1 task to the Ctx Dly DM 2 task (Fig. 5d,e). The Ctx DM 1-to-Ctx DM 2 vector reflects the difference between the gating modality 1 and the gating modality 2 processes. These results suggest that sensory, cognitive, and motor processes can be represented as vectors in the task space. Therefore, the representation of a task can potentially be expressed as a linear summation of vectors representing the underlying sensory, cognitive, and motor processes. This finding is reminiscent of previous work showing that neural networks can represent words and phrases compositionally^31^.

**Figure 5.**
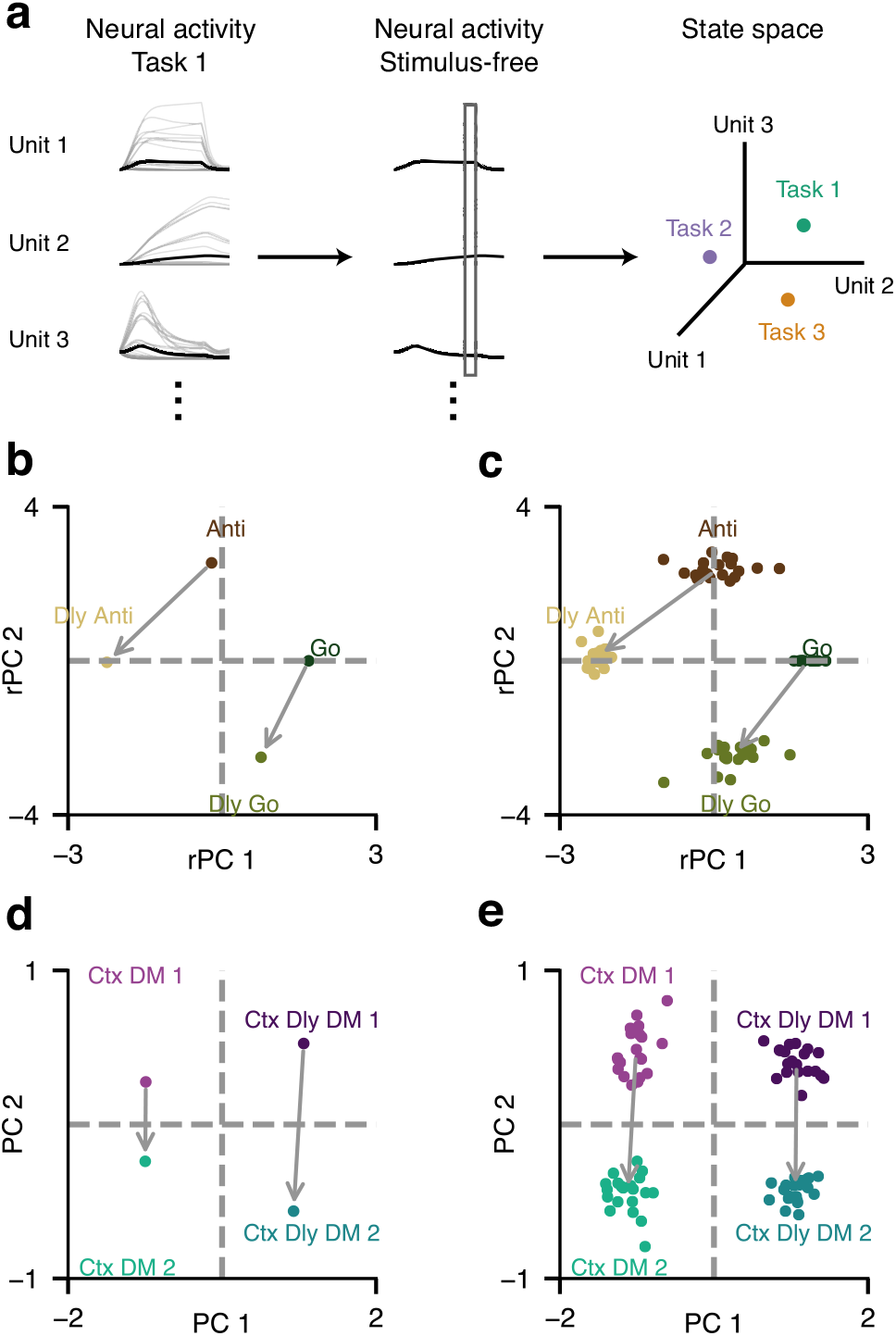
Compositional representation of tasks in state space. **(a)** The representation of each task is the population activity of the recurrent network at the end of the stimulus presentation, averaged across different stimulus conditions. **(b)** Representations of the Go, Dly Go, Anti, Dly Anti tasks in the space spanned by the top two principal components (PCs) for a sample network. For better comparison across networks, the top two PCs are rotated and reflected (rPCs) to form the two axes (see Online Methods). **(c)** The same analysis as in **(b)** is performed for 20 networks, and the results are overlaid. **(d)** Representations of the Ctx DM 1, Ctx DM 2, Ctx Dly DM 1, and Ctx Dly DM 2 tasks in the top two PCs for a sample network. **(e)** The same analysis as in **(d)** for 20 networks.

### Performing tasks with composition of rule inputs

We showed that the representation of tasks could be compositional in principle. However, it is unclear whether in our network this principle of compositionality can be extended from representing to performing tasks. The network is normally instructed which task to perform by activation of the corresponding rule input unit. What would the network do in response to a compositional rule signal as a combination of several activated and deactivated rule units? We tested whether the network can perform tasks by receiving composite rule inputs (Fig. 6a).

Consider the same two sets of tasks as in Fig. 5. The network can perform the Dly Anti task well when provided with the particular combination of rule inputs: Anti + (Dly Go - Go) (Fig. 6b). In contrast, the network fails to perform the Dly Anti task when provided with several other combinations of rule inputs (Fig. 6b). Similarly, the network can perform the Ctx Dly DM 1 task best when provided the composite rule inputs of Ctx Dly DM 2 + (Ctx DM 1 - Ctx DM 2) (Fig. 6c). In accordance with these results, we found that connection weights from individual rule input units to recurrent units also display a compositional structure (Supplementary Fig. 6). Together, these results further confirmed that our network learned the implicit compositional relationship between tasks. In such a network, learning a new task may not require any modification to the recurrent connections. Instead, it only requires learning the appropriate combination of rule inputs that control the information flow within the network^2^.

**Figure 6.**
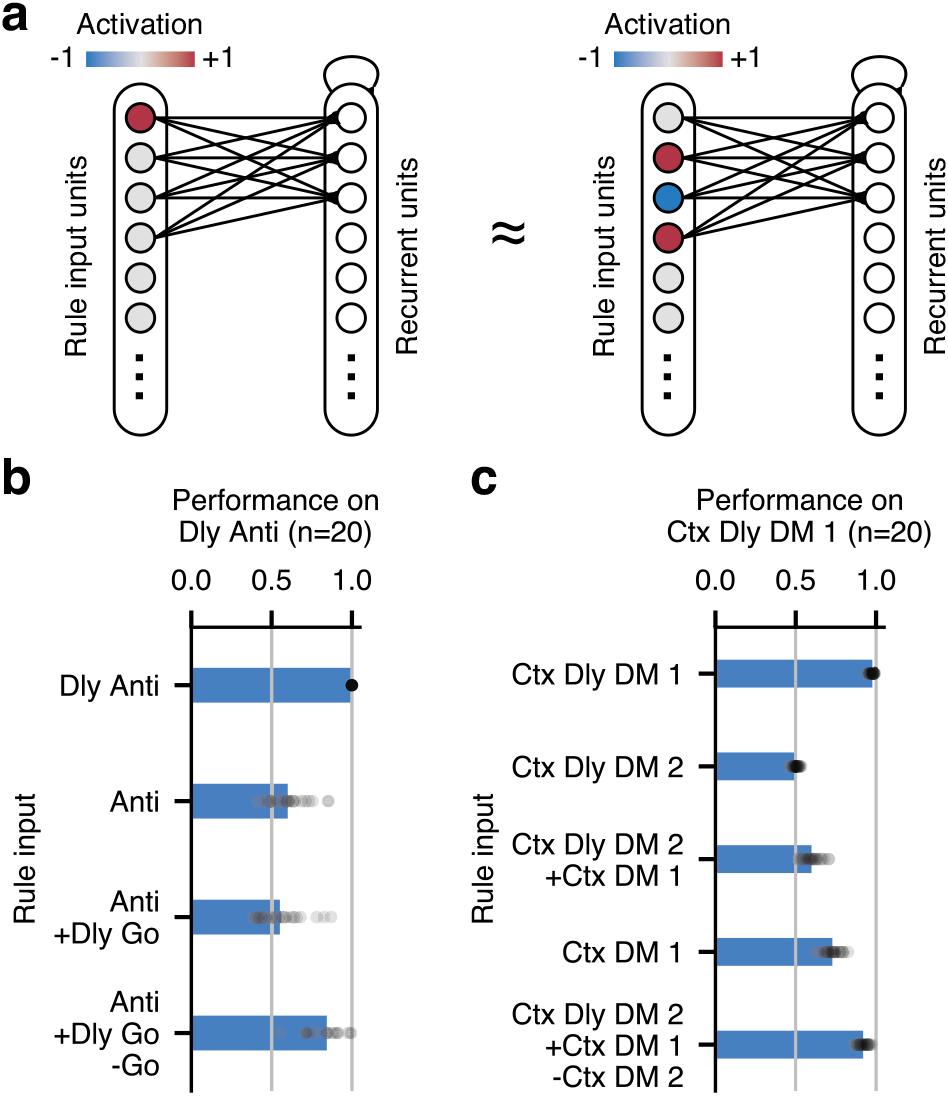
Performing tasks with algebraically composite rule inputs. **(a)** During training, a task is always instructed by activation of the corresponding rule input unit (left). After training, the network can potentially perform a task by activation or deactivation of a set of rule input units meant for other tasks (right). **(b)** The network can perform the Dly Anti task well if given the Dly Anti rule input or the Anti + (Dly Go - Go) rule input. The network fails to perform the Dly Anti task when provided other combinations of rule inputs. **(c)** Similarly, the network can perform the Ctx Dly DM 1 task well when provided with the Ctx Dly DM 2 + (Ctx DM 1 - Ctx DM 2) rule input. Circles represent results of individual networks, while bars represent median performances of 20 networks.

### Continual training of many cognitive tasks

In humans and other animals, the performance of a well-trained task can be retained, even without re-training, for months or even years. However, when using traditional network training techniques, artificial neural networks rapidly forget previously learned tasks after being exposed to new tasks. This failure of retaining memories during sequential training of tasks, termed “catastrophic forgetting,” is inevitable when using common network architectures and training methods^32,33^. Network parameters (such as connection weights) optimal for a new task can be destructive for old tasks (Fig. 7a). Recent work proposed several continual learning methods to battle catastrophic forgetting^32–34^. These methods typically involve selective protection of connection weights that are deemed important for previously learned tasks.

By employing one such technique^33^, we were able to substantially improve the performance of networks that are sequentially trained on a set of cognitive tasks (Fig. 7b). The continual learning technique is especially effective at helping the network retain performance of tasks learned earlier. For example, the network can retain high performance in a working memory task after successfully learning ten additional tasks (Fig. 7c). We analyzed the FTV distributions for three example pairs of tasks in the continual learning networks (Fig. 7d-f). The shapes of these FTV distributions can be markedly different from the corresponding ones of the interleaved-training networks (Fig. 7d,e, Fig. 4b,c). It is possible that this result depends on factors in the continual learning, such as the order of individual tasks used during training, more careful comparisons are needed in future studies. Nevertheless, our findings suggest that sequential training of tasks could drastically shape neural network representations.

## DISCUSSION

Higher-order cortical areas, especially the lateral prefrontal cortex, are remarkably versatile for their engagement in a wide gamut of cognitive functions. Here we investigated how multiple cognitive tasks are represented in a single recurrent neural network model. First, we demonstrated how the trained neural network could be dissected and understood for a family of tasks. Next, we identified clusters of units that are each specialized for a subset of tasks. Each cluster potentially represents a particular sequence of the sensori-motor events and a subset of cognitive (such as working memory, categorization, decision-making, inhibitory control) processes that are the building blocks for flexible behavior. We observed a close match between the task selectivity of each cluster and its causal role, which suggests that analysis of neural activity can provide meaningful functional insights in trained neural networks. We proposed a measure, the fractional task variance, that probes the neural relationship between a pair of tasks at the single neuron level. This measure allowed us to summarize five distinct and typical kinds of neural relationships in our network. This measure can be readily applied to firing activity of single units recorded from animals performing two or more tasks. Surprisingly, we found that the representation of tasks in our network is compositional, a critical feature for cognitive flexibility. By virtue of the compositionality, a task can be correctly instructed by composing instructions for other tasks. Finally, using a recently proposed continual learning technique, we can train the network to learn many tasks sequentially.

**Figure 7.**
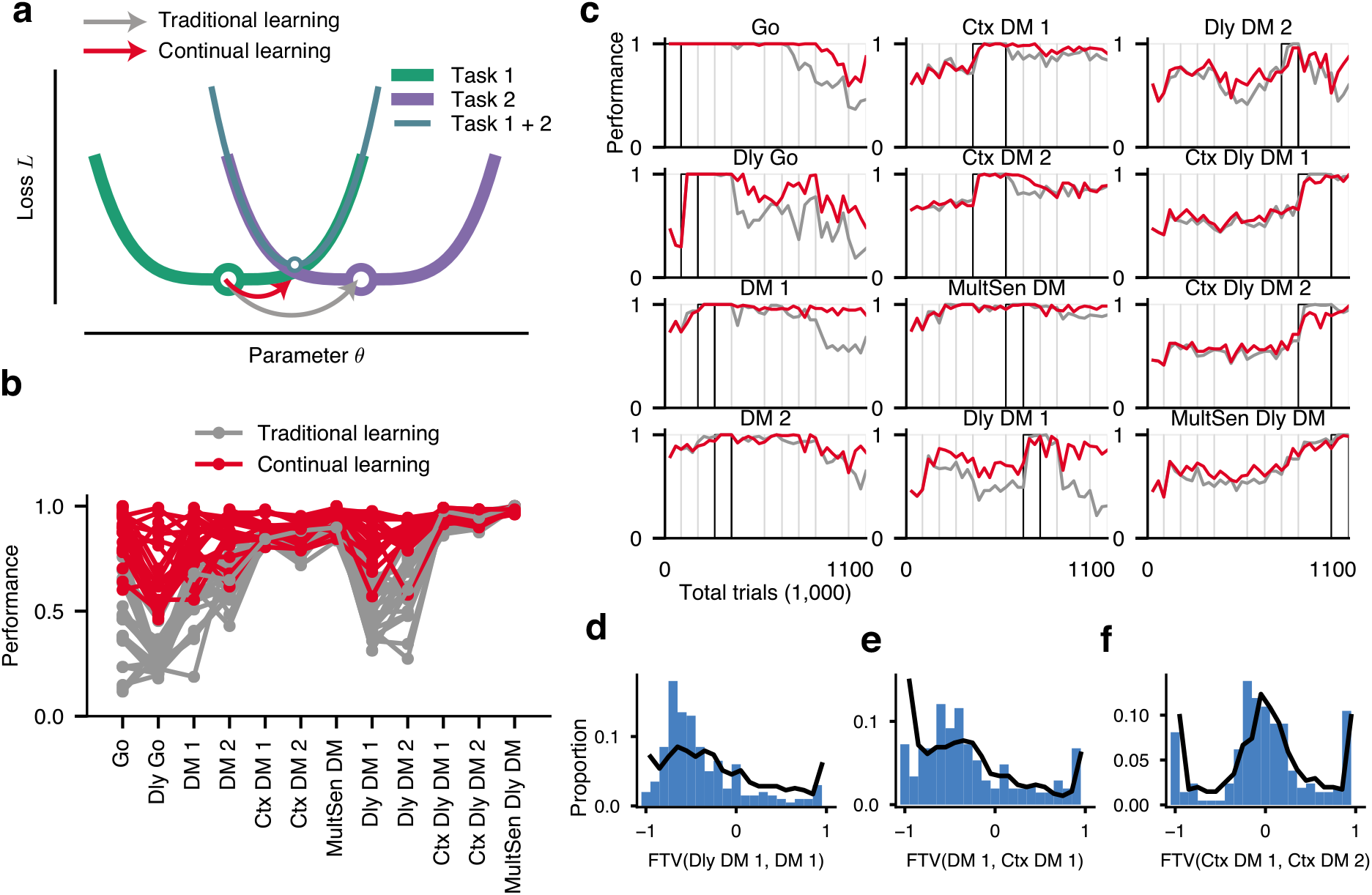
Sequential training of cognitive tasks. **(a)** Schematics of continual learning. The network learns to perform a task by modifying parameters to minimize the loss function for this task. When a network is trained on two tasks sequentially with traditional learning techniques (gray arrow), training for the second task can easily result in the failure of performing the first task, because the minima (circle) of the loss functions of tasks 1 (green) and 2 (purple) are far apart. Continual learning techniques can protect previously-learned tasks by preventing large changes of important network parameters (red arrow). Arrows show changes of an example parameter *θ* when task 2 is trained after task 1 is already learned. **(b)** Final performance across all trained tasks with traditional (gray) or continual (red) learning techniques. Only 12 tasks are trained due to difficulty of learning more tasks even with continual learning techniques. Lines represent results of individual networks. **(c)** Performance of all tasks during sequential training of one network with traditional (gray) or continual (red) learning techniques. For each task, the black box indicates the period in which this task is trained. **(d-f)** Fractional variance distributions for three pairs of tasks. Blue: distribution for one sample network. Black: averaged distribution over 20 networks.

Monkeys, and in some cases rodents, can be trained to alternate between two tasks^11,22,25^’^35^’^36^. Single-unit recordings from these experiments can potentially be analyzed to compute the fraction task variance distributions. Theoretical studies argued that for maximum cognitive flexibility, prefrontal neurons should be selective to mixtures of multiple task variables^37^. Mixed selectivity neurons are indeed ubiquitous within the prefrontal cortex^10^. We showed that most units in our network are strongly selective to rules (Fig. 3). Meanwhile, these units are selective to other aspects of tasks (otherwise their task variances would be zero). For example, the Anti units (Fig. 2) are highly activated only during the Anti tasks and when the stimulus is in their preferred directions. Therefore, units in our network display strong nonlinear mixed selectivity, as found in neurons of the prefrontal cortex^10^. Conceptually, this work extends the notion of mixed selectivity from within a single task to across multiple tasks.

Multiple cognitive tasks are more common in human imaging studies. In a series of experiments, Cole and colleagues trained humans to perform 64 cognitive tasks following compositional rule instructions^6,38^. They trained linear classifiers to decode rules from prefrontal neural activity patterns. These classifiers can significantly generalize to novel tasks^6^, consistent with a compositional neural representation of rules. Although trained with discrete rule instructions, our network develops a clear compositional structure in its representations, as shown using the population activity at a single time point (near the end of stimulus presentation). Temporal dynamics in neural circuits are ubiquitous during cognitive tasks^39^ and are potentially critical for cognitive computations^18^, so the study of steady-state responses here is merely a first step towards understanding the dynamical representation of tasks. Future work could study how dynamical representations of tasks are related to one another in the state space. Cole et al. found that humans can rapidly adapt to new tasks by adjusting the functional connectivity patterns of parietal-frontal flexible hubs^38^. In the future, graph-theoretic analysis can be used to test whether our trained network developed flexible hubs that coordinate information flow across the network. There exists a structural hierarchy within the human prefrontal cortex, with more abstract cognitive processes being represented in the more anterior areas^40,41^. It is unclear if our trained network developed hierarchical representations of cognitive processes or tasks. If it did, a subset of units should represent more abstract aspects of the tasks, while other units represent the concrete, sensorimotor aspects. This question is hard to address for now because the 20 tasks we chose are not organized in a clearly hierarchical way^40^.

Training artificial neural networks for multiple tasks has a long history in the field of machine learning^42^. However, it has mainly been used as a method to improve training and generalization. There were few studies on the representation of task structure or task set in trained networks. Modern artificial neural networks are capable of highly complex tasks, such as playing Atari games^43^, which likely involve a range of cognitive skills. However, in contrast to cognitive tasks that are specifically designed to shed light on neural mechanisms of cognition, complex real-life tasks remain challenging to analyze. In principle, we can strike a balance between the two approaches by designing a set of tasks that are complex enough, yet still amenable to analysis. The ability to “open the box” and elucidate the inner working of the network after training is crucial for understanding neural mechanisms of cognition in neuroscience.

Like other works on trained neural networks^11,14–19,44^, the machine learning protocol we used is not validated biologically. Besides, our RNN consists of a single neural population, in contrast to the brain system where a number of interacting brain regions are engaged in a cognitive task^22,45^. Although our neural network model developed functionally specialized clusters of units through training, it is unclear how to map them onto different brain areas. Furthermore, in our network, a rule input is explicitly provided throughout the trial, therefore there is no need for the network to hold the “task set” internally using persistent activity^4,5^. This, however, can be remedied by providing the rule cue only at the beginning of each trial, which would encourage the network to internally sustain the task set. We can even ask the network to figure out a task rule by trial-and-error^46^. In spite of these concerns, our approach offers an efficient computational platform to test hypotheses about neural representations and mechanisms that could guide experiments and data analysis. Furthermore, this approach can yield new conceptual insights, as shown here by the finding of compositional task representation. Future progress in this direction, at the interface between neuroscience and artificial intelligence, will advance our understanding of flexible behavior in many cognitive tasks.

## ACKNOWLEDGMENTS

We thank current and former members of the Wang lab, especially S.Y. Li, O. Marschall, and M. Joglekar, and E. Ohran for fruitful discussions; J.A. Li, J.D. Murray, D. Ehrlich, and J. Jaramillo for critical comments on the manuscript; and S. Wang for assistance with the NYU HPC clusters. This work was supported by an Office of Naval Research Grant N00014-13-1-0297, a National Science Foundation Grant Number 16-31586, and a Google Computational Neuroscience Grant (X.J.W.) and a Samuel J. and Joan B. Williamson Fellowship (G.R.Y.).

## AUTHOR CONTRIBUTIONS

G.R.Y. and X.J.W. designed the study. G.R.Y., H.F.S, W.T.N, and X.J.W. had frequent discussions. G.R.Y. performed the research. G.R.Y., H.F.S, W.T.N, and X.J.W. wrote the manuscript.

## COMPETING FINANCIAL INTERESTS

The authors declare no competing financial interests.

## DATA AVAILABILITY STATEMENT

All codes will be available at publication.

## ONLINE METHODS

### Code availability

All codes are available on GitHub.

### Network structure

The recurrent neural networks shown in the main text all contain *N_rec_* = 256 units. The results are largely insensitive to the network size. Similar results were obtained in networks of sizes between 128 and 512 units (the range we tested). The network is a time-discretized recurrent neural network with positive activity^15^. Before time-discretization, the network activity r follows a continuous dynamical equation

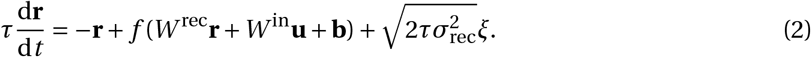

In this equation, **u** is the input to the network, **b** is the bias or background input, τ = 100ms is the neuronal time constant, *f* (·) is the neuronal nonlinearity that keeps the unit activity nonnegative, *ξ* are *N_rec_* independent Gaussian white noise processes with zero mean and unit variance, and σ_rec_ = 0.05 is the strength of the noise. In particular, we use a standard softplus function

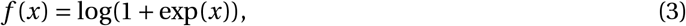

which after re-parameterization is very similar to a neuronal nonlinearity, i.e., f-I curve, commonly used in previous neural circuit modelings^28^. A set of output units **z** read out nonlinearly from the network,

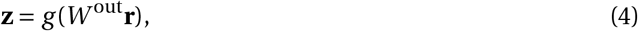

where *g*(*x*) = 1/(1 + exp(*-x*)) is the logistic function, bounding output activities between 0 and 1. *W*^in^, *W*^rec^, *W*^out^ are the input, recurrent, and output connection matrices respectively.

After using the first-order Euler approximation with a time discretization step Δ*t*, we have

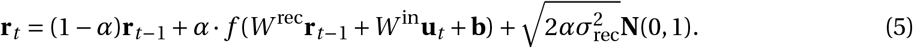

Here *α* ≡ Δ*t*/τ, and **N**(0,1) stands for the standard normal distribution. We use a discretization step Δ*t* = 20ms. We imposed no constraint on the sign or the structure of the weight matrices *W*^in^, *W*^rec^, *W*^out^. The network and the training are implemented in TensorFlow^47^.

The network receives four types of noisy inputs,

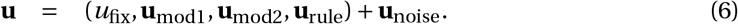

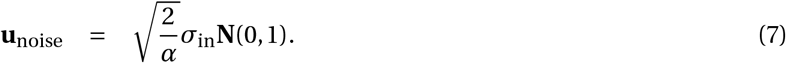

Here the input noise strength **N**(0,1) = 0.01. The fixation input *u*_fix_ is typically at the high value of 1 when the network should fixate. The fixation input goes to zero when the network is required to respond. The stimulus inputs **u**_mod1_ and **u**_mod2_ comprise two “rings” of units, each representing a one-dimensional circular variable described by the degree around a circle. Each ring contains 32 units, whose preferred directions are uniformly spaced from 0 to 2π. For unit *i* with a preferred direction *θ_i_*, its activity for a stimulus presented at direction *Ψ* is

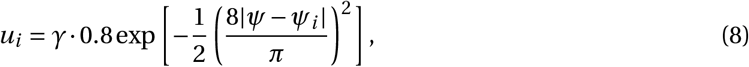

where *γ* is the strength of the stimulus. For multiple stimuli, input activities are added together. The network also receives a set of rule inputs **u**_rule_ that encode which task the network is supposed to perform on each trial. Normally, **u**_rule_ is a one-hot vector. That means the rule input unit corresponding to the current task is activated at 1, while other rule input units remain at 0. Therefore the number of rule input units equals to the number of tasks trained. For compositional rule inputs (Fig. 6), the activation of rule input units can be an arbitrary pattern. For example, for the combined rule input Anti + (Dly Go - Go), the activities of the rule input units corresponding to the Go, Dly Go, and Anti tasks are −1, +1, and +1 respectively. In total there are *N*_in_ = 1 + 32 × 2 + 20 = 85 input units.

The network projects to an output ring zout, which also contains 32 units. The output ring units encode the response directions using similar tuning curves to the ones used for the input rings. In addition, the network projects to a fixation output unit **z**_fix_, which should be at the high activity value of 1 before the response and at 0 once a response is generated. In total there are *N*_out_ = 1 + 32 = 33 output units.

We lesion a network unit by setting to zero its projection weights to all recurrent and output units.

### Tasks and performances

Here we first describe the common setup for the 20 tasks trained. Deviations from the common setup will be described below individually. The rule input unit corresponding to the current task will be activated throughout the whole trial. The network receives a fixation input, which is activated from the beginning of the trial. When the fixation input is on, the network should fixate by having the fixation output unit at a high activity **ẑ**_fix_. The offset of the fixation input usually indicates the onset of the response or go epoch, when the network needs to report the response direction through activities of the output ring. During the response epoch, the fixation output unit has a target output of **ẑ**_fix_ = 0.05. For a target response direction Ψ, the target output activity of an output unit *i* is

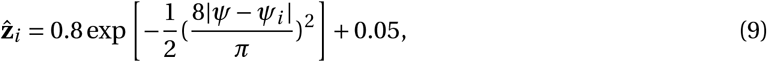

where *Ψ_i_* is the preferred response direction of unit *i*. When no response is required, the target output activity is fixed at **ẑ**_*i*_ = 0.05. The network also receives one or two stimuli. Each stimulus contains information from modality 1, 2, or both. When there is only one stimulus, the direction of the stimulus is drawn from a uniform distribution between 0 and 360 degree.

A trial is considered correct only if the network correctly maintained fixation and responded to the correct direction. The response direction of the network is read out using a population vector method. The decoded response direction is considered correct if it is within 36 degrees of the target direction. If the activity of the fixation output falls below 0.5, the network is considered to have broken fixation.

The discrimination thresholds *a* in Supplementary Fig. 2 are obtained by fitting Weibull functions to performances *p* as a function of coherences *c* at a fixed stimulus duration,

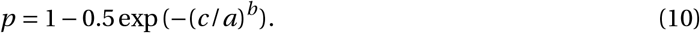

Each task can be separated into distinct epochs. Fixation (fix) epoch is the period before any stimulus is shown. It is followed by the stimulus epoch 1 (stim1). If there are two stimuli separated in time, then the period between the two stimuli is the delay epoch, and the second stimulus is shown in the stimulus epoch 2 (stim2). The period when the network should respond is the go epoch. The duration of the fixation, stim1, delay1, stim2, and go epochs are *T*_fix_, *T*_stim1_, *T*_delay1_, *T*_stim2_, *T*_go_ respectively. For convenience, we grouped the 20 tasks into five task families: the Go, Anti, Decision-Making (DM), Delayed Decision-Making (Dly DM), and Matching families.

#### Go task family

This family of tasks includes the Go, RT Go, and Dly Go tasks. In all three tasks, a single stimulus is randomly shown in either modality 1 or 2, and the response should be made in the direction of the stimulus. These three tasks differ in their stimulus onset and offset times. In the Go task, the stimulus appears before the fixation cue goes off. In the RT Go task, the fixation input never goes off, and the network should respond as soon as the stimulus appears. In the Dly Go task, a stimulus appears briefly and is followed by a delay period until the fixation cue goes off. The Dly Go task is similar to the memory-guided saccade task^20^.

For the Go task,

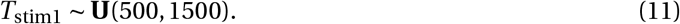

**U**(*t*_1_, *t*_2_) is a uniform distribution between *t*_1_ and *t*_2_. The unit for time is ms and is omitted for brevity. For the RT Go task,

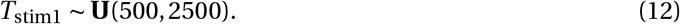

For the Dly Go tasks,

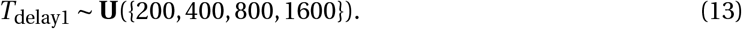

Here **U**({*a*_1_,…, *a*_n_}) denotes a discrete uniform distribution over the set {*a*_1_,…, *a*_n_}.

#### Anti task family

This family includes the Anti, RT Anti, and Dly Anti tasks. These three tasks ł are the same as their counterpart Go-family tasks, except that the response should be made to the opposite direction of the stimulus.

#### DM family

This family includes five perceptual decision making tasks: the DM 1, DM 2, Ctx DM 1, Ctx DM 2, and MultSen DM tasks. In each trial, two stimuli are shown simultaneously and are presented till the end of the trial. Stimulus 1 is drawn randomly between 0 and 360 degree, while stimulus 2 is drawn uniformly between 90 and 270 degree away from stimulus 1. In DM 1, the two stimuli only appear in modality 1, while in DM 2, the two stimuli only appear in modality 2. In DM 1 and DM 2, the correct response should be made to the direction of the stronger stimulus (the stimulus with higher γ). In Ctx DM 1, Ctx DM 2, and MultSen DM tasks, each stimulus appears in both modality 1 and 2. In the Ctx DM 1 task, information from modality 2 should be ignored, and the correct response should be made to the stronger stimulus in modality 1. In the Ctx DM 2 task, information from modality 1 should be ignored. In the MultSen DM task, the correct response should be made to the stimulus that has a stronger combined strength in modalities 1 and 2.

The DM 1 and DM 2 tasks are inspired from classical perceptual decision making tasks based on random-dot motion stimuli^21^. In random-dot motion tasks, there is only one stimulus, the coherence of which is varied across trials. Following the tradition of Wang, 2002^30^, we use two input stimuli to model momentary motion evidence towards the two target directions. When the two stimuli have the same strengths (γ_1_ = γ_2_), there is no net evidence towards any target direction, mimicking the condition of 0 motion coherence in the random-dot motion task. A stronger difference in the stimulus strengths emulates a stronger motion coherence. For a coherence *c* representing net evidence for the direction of stimulus 1, the strengths of stimulus 1 and 2 (γ_1_, γ_2_) are set as

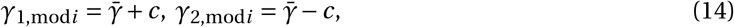

respectively, where *i ϵ* 1,2 is the modality. Here 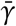 is the average strength of the two stimuli. For each trial, we draw 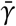 from a uniform distribution around 1, *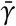 ~* **U**(0.8,1.2). Indeed, in all DM-family tasks and Dly DM-family tasks, there is a single coherence *c* in each trial that determines the overall strength of net evidence towards the direction represented by stimulus 1. For all DM family tasks,

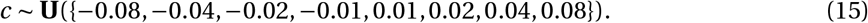

The duration of stimulus 1, which is fixed in each trial, is drawn from the following distribution,

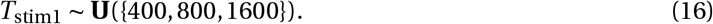

Indeed, all tasks from the DM family use the same distribution for *T*_stim1_. And since the two is stimuli are shown simultaneously, *T*_stim1_ = *T*_stim2_.

The Ctx DM 1 and Ctx DM 2 tasks are inspired from context-dependent decision-making tasks performed by macaque monkeys^11^. Now each stimulus is presented in both modalities at the same direction, with strengths γ_1, mod1_, γ_1, mod2_ for stimulus 1, and γ_2, mod1_, γ_2, mod2_ for stimulus 2. The stimulus strengths are determined by the coherence for modality 1 and 2 (*c*_mod1_, *c*_mod2_), so we have

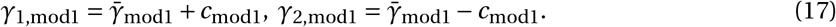

Similar equation holds for modality 2 as well. *c*_mod1_ and *c*_mod2_ are drawn independently from the same distribution. In Ctx DM 1, *c* = *c*_mod1_, while in Ctx DM 2, 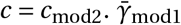 and 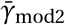 are also drawn from **U**(0.8,1.2). In the original Mante task^11^, there is an additional delay period between the stimuli and the response period, which is not included here.

The MultSen DM task mimics a multi-sensory integration task^23^. The setup of stimulus is similar to those in the Ctx DM 1 and Ctx DM 2 tasks, except that the network should integrate information from both modalities and the stronger stimulus is the one with higher averaged strength from modality 1 and 2. The overall coherence *c* = (*c*_mod1_ + *c*_mod2_)/2. We determine all four strengths with the following procedure. First we determine the average strength of stimulus 1 across both modalities, γ_1_, and the average strength of stimulus 2, γ_2_.

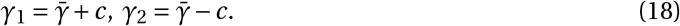

Here 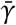 and *c* both follow the same distributions as other DM-family tasks. Then we set

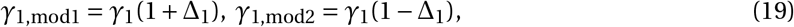

where Δ_1_ ~ **U**(0.1,0.4)∪**U**(-0.4,-0.1). Similarly for stimulus 2.

#### Dly DM family

This family includes Dly DM 1, Dly DM 2, Ctx Dly DM 1, Ctx Dly DM 2. These ł tasks are similar to the corresponding tasks in the DM family, except that in the Dly DM family tasks, the two stimuli are separated in time. The Dly DM 1 and Dly DM 2 tasks are inspired by the classical parametric working memory task developed by Romo and colleagues^24^. The two stimuli are both shown briefly and are separated by a delay period. Another short delay period follows the offset of the second stimulus.

For all Dly DM family tasks,

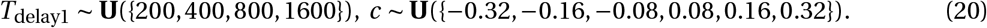

And, *T*_stim1_ = *T*_stim2_ = 300.

#### Matching family

This family of tasks includes the DMS, DNMS, DMC, DNMC tasks. In these tasks, two stimuli are presented consecutively and separated by a delay period. Each stimulus can appear in either modality 1 or 2. The network response depends on whether or not the two stimuli are “matched." In the DMS and DNMS tasks, two stimuli are matched if they point towards the same direction, regardless of their modalities. In DMC and DNMC tasks, two stimuli are matched if their directions belong to the same category. The first category ranges from 0 to 180 degrees, while the rest from 180 to 360 degrees belongs to the second category. In the DMS and DMC tasks, the network should respond towards the direction of the second stimulus if the two stimuli are matched and maintain fixation otherwise. In the DNMS and DNMC tasks, the network should respond only if the two stimuli are not matched, i.e., a non-match, and fixate when it is a match.

During training of these tasks, half of the trials are matching, and the other half are non-matching. In DMS and DNMS tasks, stimulus 1 is always drawn randomly. In half of the trials, stimulus 2 appears at the same direction as stimulus 1. In the other half, stimulus 2 is drawn randomly between 10 and 350 degree away from stimulus 1. In DMC and DNMC tasks, both stimulus 1 and 2 are drawn randomly and independently from the uniform distribution

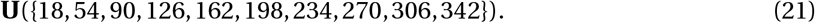

In all Matching family tasks,

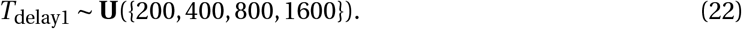

Also, match trials and non-match trials always appear with equal probability.

### Training procedure

The loss *ℒ* to be minimized is computed by time-averaging the squared errors between the network output **z**(*t*) and the target output **ẑ**(*t*).

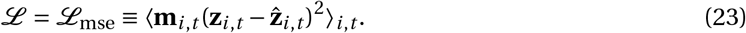

Here *i* is the index of the output units. The squared errors at different time points and of different output units are potentially weighted differently according to the non-negative mask matrix **m**_i,t_. For the output ring units, before the response epoch, we have **m**_*i,t*_ = 1. The first 100ms of the response epoch is a grace period with **m**_*i,t*_ = 0, while for the rest of the response epoch, **m**_*i,t*_ = 5. For the fixation output unit, **m**_*i,t*_ is two times stronger than the mask for the output ring units.

The training is performed with Adam, a powerful variant of stochastic gradient descent^48^. We used the default set of parameters. The learning rate is 0.001, the decay rate for the 1st and 2nd moment estimates are 0.9 and 0.999 respectively.

The recurrent connection matrix is initialized with a scaled identity matrix *q ·* **1**^49^, where **1** is the identity matrix. We chose *q* = 0.54 such that the gradient is roughly preserved during backpropagation when the network is initialized. The input and output connection weights are initialized as independent Gaussian random variables with mean 0, and standard deviations 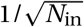 and 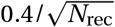 respectively. The standard deviation value for the output weights is chosen to prevent saturation of output units after initialization.

During training, we randomly interleaved all the tasks with equal probabilities, except for the Ctx DM 1 and Ctx DM 2 tasks that appear five times more frequently, because without sufficient training, the network gets stuck at an alternative strategy. Instead of correctly ignoring modality 1 or 2, the network can choose to ignore the context and integrate information from both modalities equally. This strategy gives the network an accuracy close to 75%. During training, we used mini-batches of 64 trials, in which all trials are generated from the same task for computational efficiency.

### Analysis of the Anti task family

Anti units in Fig. 2 are defined as those units that have higher summed task variance (see next section for definition) for the Anti family of tasks (S_Anti_ =Anti, RT Anti, Dly Anti) than for all other tasks. So a unit *i* is an Anti unit if

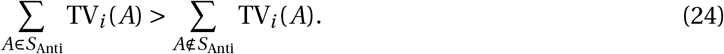

### Task variance analysis

A central goal of our analysis was to determine whether individual units within the network are selective to different tasks, or whether units tended to be similarly selective to all tasks. To quantify how selective a unit is in one task, we defined a task variance metric. To compute the task variance TV*_i_(A)* for task A and unit *i*, we ran the network for many stimulus conditions that span the space of possible stimuli. For example, in the DM family tasks, we ran the network for stimuli with directions ranging from 0 to 360 degrees and with coherences ranging from almost 0 to 0.2. After running the network for many stimulus conditions, we computed the variance across stimulus conditions (trials) at each time point for a specific unit then averaged the variance across all time points to get the final task variance for this unit. The fixation epoch is excluded from this analysis. This process was repeated for each unit in the network. Therefore

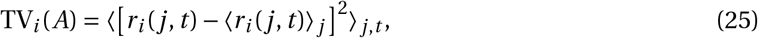

where *r_i_* (*j, t*) is the activity of unit *i* on time *t* of trial *j*. In Fig. 2,3,4, we only analyzed active units, defined as those that have summed task variance across tasks higher than a threshold, 10^−3^. The results do not depend strongly on the choice of the threshold. This procedure prevents units with extremely low task variance from being included in the analysis.

By computing each unit’s selectivity across different stimulus conditions, we naturally include the selectivity to motor outputs, because motor outputs depend ultimately on the stimuli. A unit that is only selective to motor outputs or other cognitive variables in a task will still have a non-zero task variance. Units that are purely selective to rules and/or time will, however, have zero task variance and therefore be excluded from our analysis.

The clustering of units based on their task variance patterns in Fig. 3 uses K-means clustering from the Python package scikit-learn. To assess how well a clustering configuration is, we computed its silhouette coefficient based on intra-cluster and inter-cluster distances. A higher silhouette coefficient means a better clustering. The optimal number of clusters 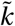 is determined by choosing the first *k* such that the silhouette coefficient for *k* + 1 clusters is worse than *k* clusters.

In Fig. 3d, we visualize the clustering using t-distributed Stochastic Neighbor Embedding (tSNE). For each unit, the normalized task variances across all tasks form a 20 dimensional vector that is then embedded in a 2-dimensional space. For the tSNE method, we used the exact method for gradient calculation, a learning rate of 100, and a perplexity of 30.

The fractional task variance with respect to tasks A and B is

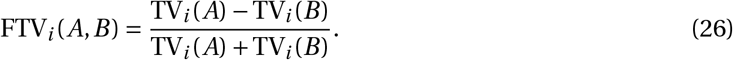

To obtain a statistical baseline for the FTV distributions as in Supplementary Fig. 4, we transform the neural activities of the network with a random orthogonal matrix before computing the task variance. For each network, we generate a random orthogonal matrix *M* using the Python package Scipy. All network activities are multiplied by this matrix *M* to obtain a rotated version of the original neural representation.

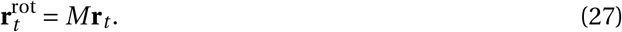

Since multiplying neural activities by an orthogonal matrix is equivalent to rotating the neural representation in state space, this procedure will preserve results from state space analysis. We then compute task variances and fractional task variances using the rotated neural activities. The FTV distributions using the rotated activities are clearly different from the original FTV distributions.

### State-space analysis

To compute the representation of a task in the state space, we first computed the neural activities across all possible stimulus conditions, then we averaged across all these conditions. For simplicity of the analysis, we chose to analyze only the steady state responses during the stimulus epoch. We do so by focusing on the last time point of the stimulus epoch, *t*_stim1,end_. So the representation of task A is

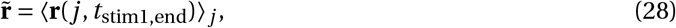

where **r**(*j*, *t*) is the vector of network activities at trial *j* and time *t* during task *A*.

For each set of tasks, we performed principal component analysis to get the lower dimensional representation. We repeated this process for different networks. The representations of Go, Anti, Dly Go, and Dly Anti tasks are close to four vertices of a square. As a result, the top two principal components have similar eigenvalues and are therefore interchangeable. To better compare across networks in Fig. 5b,c, we allowed a rotation and a reflection within the space spanned by the top two PCs. For each network, the rotated and reflected PCs (rPCs) are chosen such that the Go task representation lies on the positive part of the x-axis, and the Dly Go task lies below the x-axis. The representation of Ctx DM 1, Ctx DM 2, Ctx Dly DM 1, and Ctx Dly DM 2 tasks do not form a square, so we only allowed reflections such that Ctx Dly DM 1 is in the first quadrant. The reflected PCs are still PCs.

### Continual learning

For continual learning in Fig. 7, tasks appear sequentially. Each task is trained for 150,000 trials. Ctx DM 1 and Ctx DM 2 are still trained together and interleaved, and so are Ctx Dly DM 1 and Ctx Dly DM 2. We added a regularizer that protects old tasks by setting an additional penalty for deviations of important synaptic weights (or other parameters)^33^. When training the *μ*-th task, the regularizer is

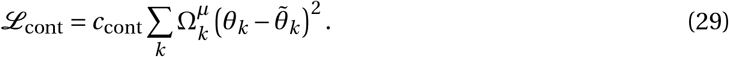

Here *c*_cont_ is the overall strength of the regularizer, *θ_k_* denotes the *k*-th parameter of the network.

The value of the anchor parameter 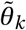*_k_* is the value of *θ_k_* at the end of the last task (the (*μ* − 1)-th task). No regularizer is used when training the first task. And 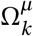 measures how important the parameter is. Notice that two recent proposals^32,33^ for continual learning both use regularizers of this form. The two proposals differ only in how the synaptic importances are computed. We chose the method of Zenke et al. 2017, because the method of Kirkpatrick et al. 2017 measures the synaptic importance locally in the parameter space, resulting in underestimated and inaccurate synaptic importance values for our settings. In Zenke et al. 2017, the importance of one parameter is determined using this parameter’s historic contribution to the change in the loss function. For the *k*-th parameter, the contribution to the change in loss during task *μ* is

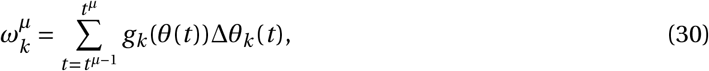

where *g*_*k*_ (θ(*t*)) is the gradient of loss with respect to *θ_k_* evaluated at *θ*_*k*_(*t*), i.e., 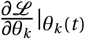, and *Δθ_k_(t)* is the parameter change taken at step *t*. Therefore 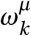 tracks how parameter *θ_k_* contributes to changes in the loss during the *μ*-th task (from *t^μ-1^* to *t*^μ^). The final synaptic importance is computed by first normalizing 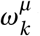 with the total change in the synaptic weight 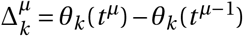, and summing 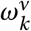 for all tasks *v* < μ.

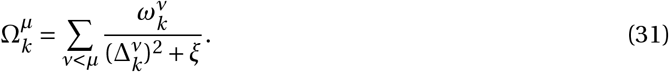

The additional hyperparameter *ξ* prevents 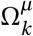 from becoming too large. The hyperparameters *c* = 0.1 and *ξ* = 0.01 are determined by a coarse grid search. The final loss is the sum of the squared-error loss and the continual learning regularizer.

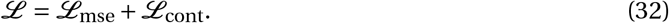

Even with the help of the continual learning technique, we had difficulties training the network using our original task setups. So we made the decision-making tasks easier by increasing the coherences by 4 times. We also made the delayed decision-making tasks easier by increasing the coherence by 2 times. In addition, we used the rectified linear function as the neuronal non-linearity, namely *f* (*x*) = max(*x*,0). We found that networks using rectified linear units learned context-dependent tasks (Ctx DM 1, Ctx DM 2, Ctx Dly DM 1, and Ctx Dly DM 2) more easily.

---

**Supplementary Fig. 1.**
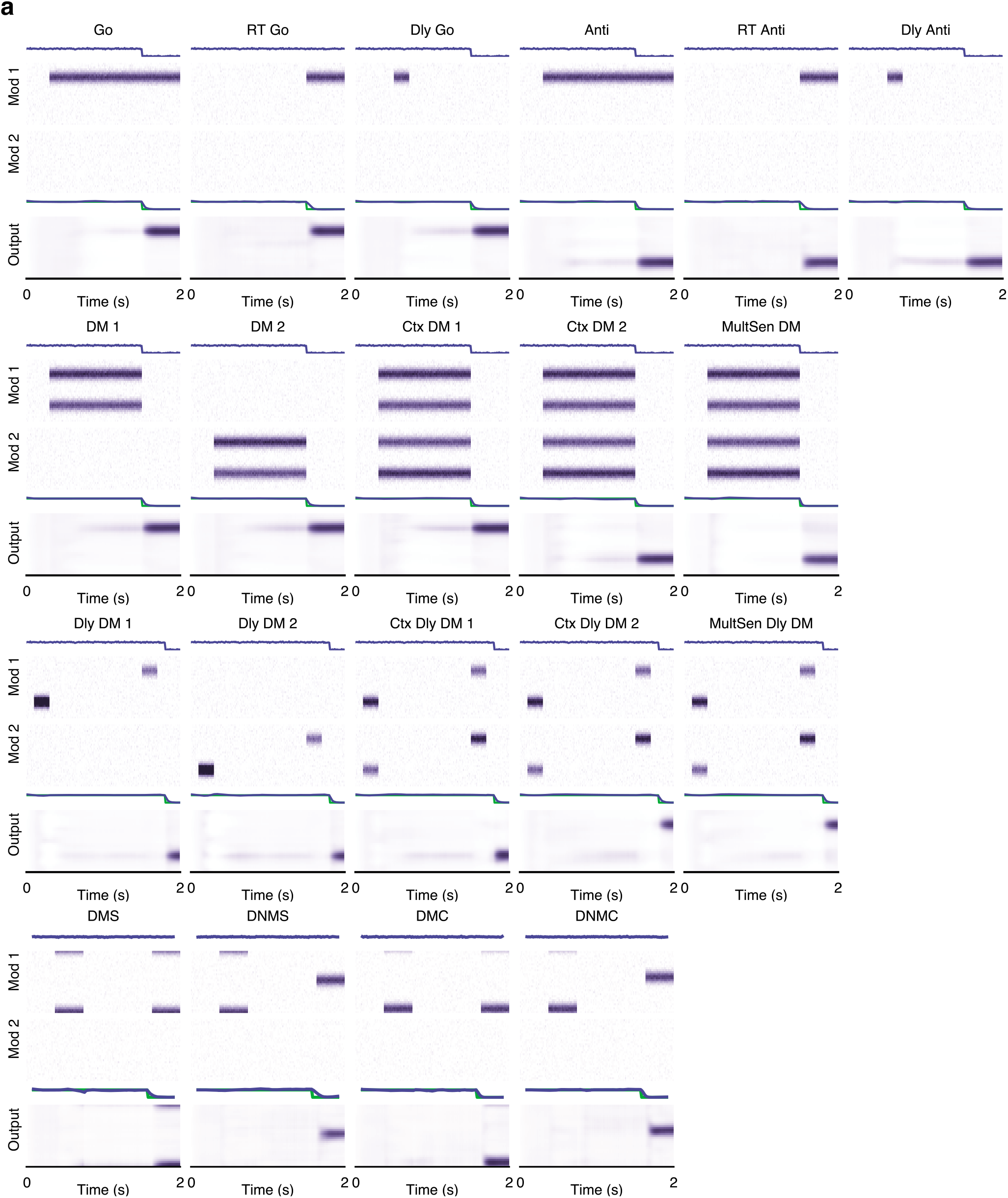
Sample trials from the 20 tasks trained. **(a)** Convention is the same as Fig. 1a. Output activities are obtained from a sample network after training. Green lines are the target activities for the fixation output unit.

**Supplementary Fig. 2.**
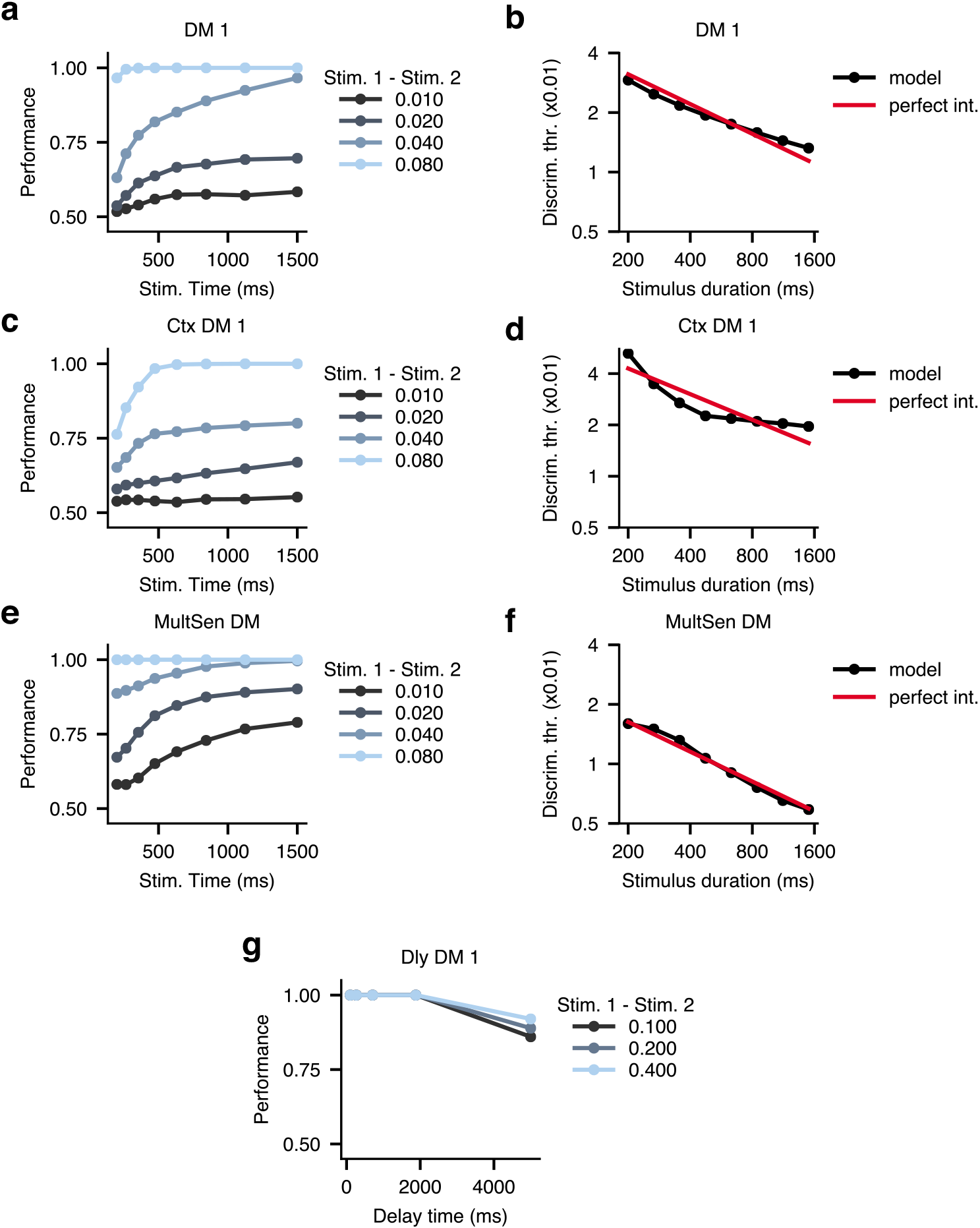
Psychometric tests for a range of tasks. **(a)** Decision making performances improve with longer stimulus presentation time and stronger stimulus coherence in the DM 1 task in a sample network. **(b)** Discrimination thresholds decrease with longer stimulus presentation time in the DM 1 task. The discrimination thresholds are estimated by fitting cumulative Weibull functions. **(c-f)** Same analyses as **(a,b)** for the Ctx DM 1 **(c,d)** and MultSen DM **(e,f)** task. These results are obtained from one sample network. The capability to integrate information over time varies across networks. However, this variation has no impact on other results. **(g)** A sample network is able to perform well above chance in the Dly DM 1 task for a delay period of up to five seconds.

**Supplementary Fig. 3.**
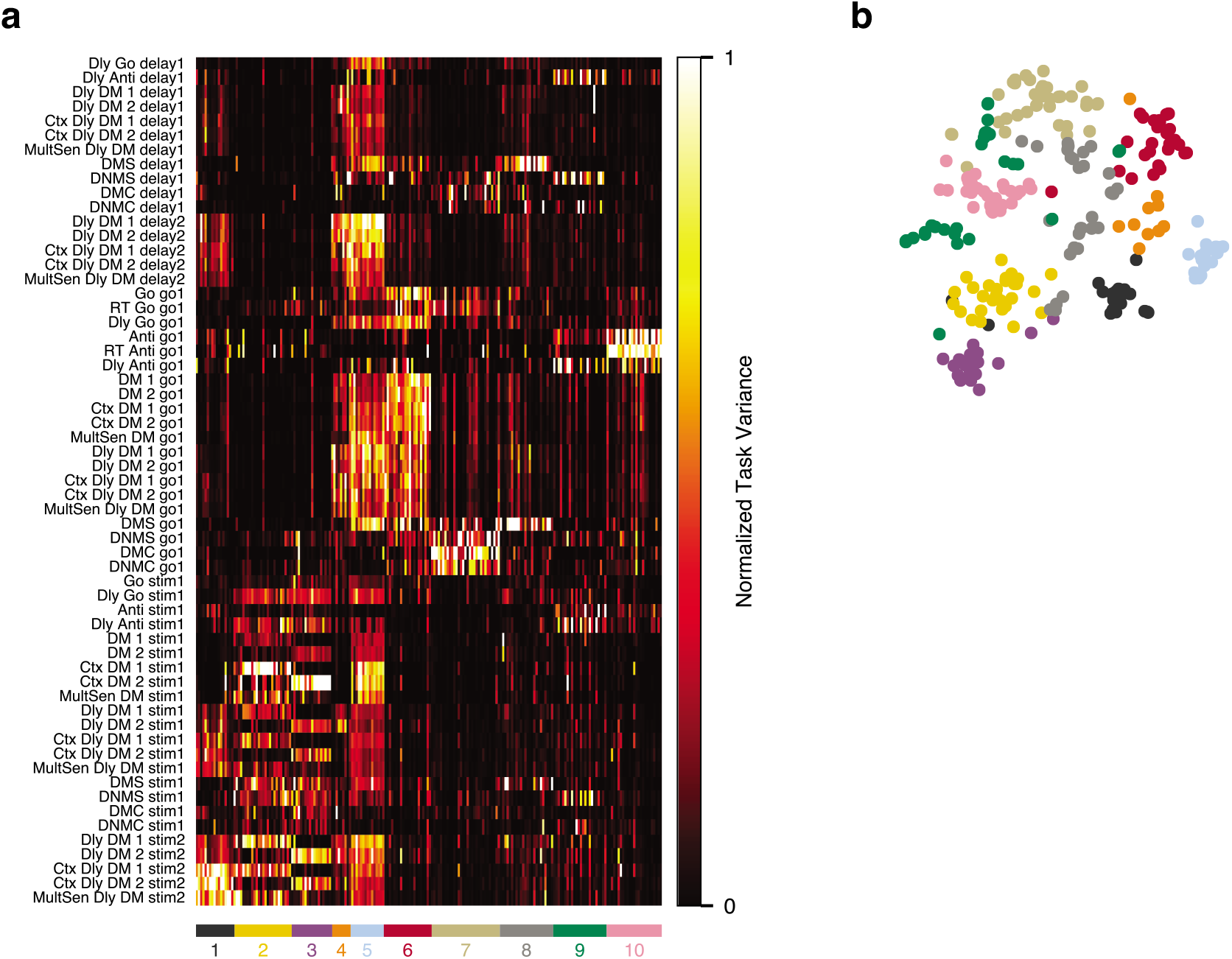
Epoch variances across all task epochs and active units. **(a)** Epoch variance is computed in a similar way to task variance, except that it is computed for individual task epochs instead of tasks. There are clusters of units that are selective in specific epochs. **(b)** Visualization of the epoch variance map in the same style as Fig. 3d.

**Supplementary Fig. 4.**
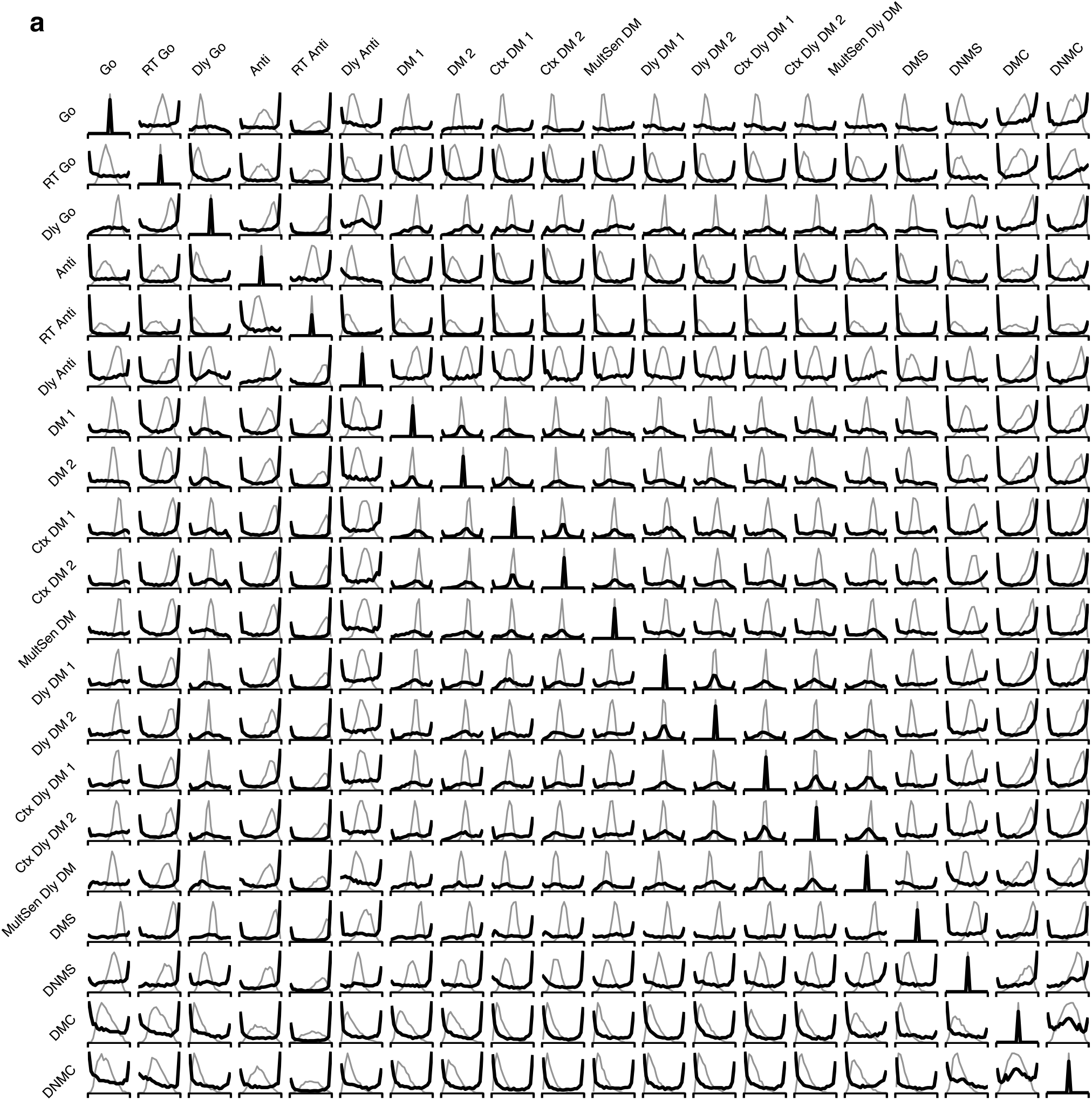
Fractional variance distributions for all pairs of tasks. **(a)** There is a total of 190 unique pairs of tasks from all 20 tasks trained. Each fractional variance distribution (black) shown here is averaged across 20 networks. As a control, we also computed fractional variance distributions (gray) from activities of surrogate units that are generated by randomly mixing activities of the original network units (see Online Methods).

**Supplementary Fig. 5.**
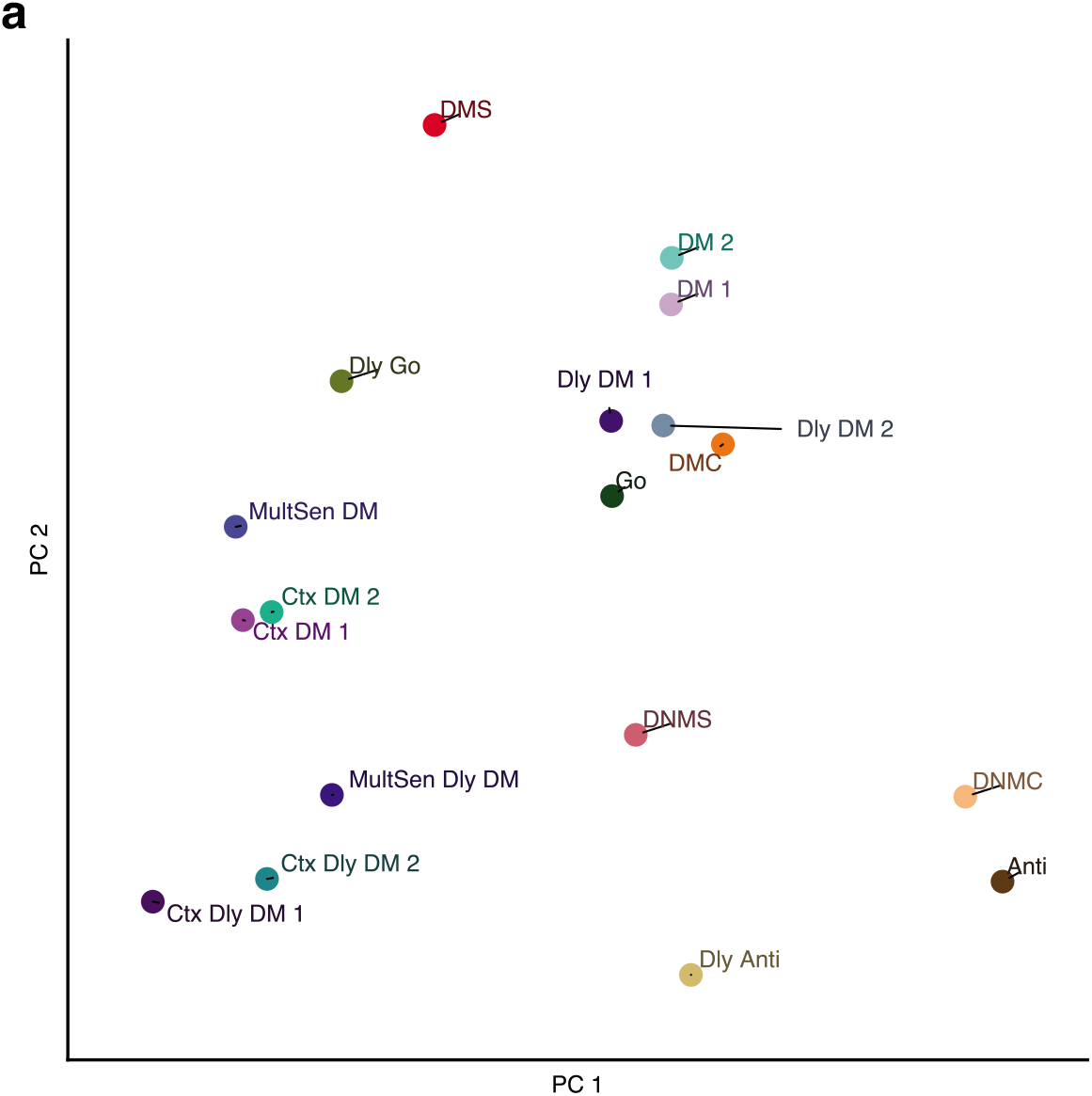
Representation of all tasks in state space. **(a)** The representation of each task is computed the same way as in Fig. 5. Here showing the representation of all tasks in the top two principal components. RT Go and RT Anti tasks are not shown here because there is no well-defined stimulus epoch in these tasks.

**Supplementary Fig. 6.**
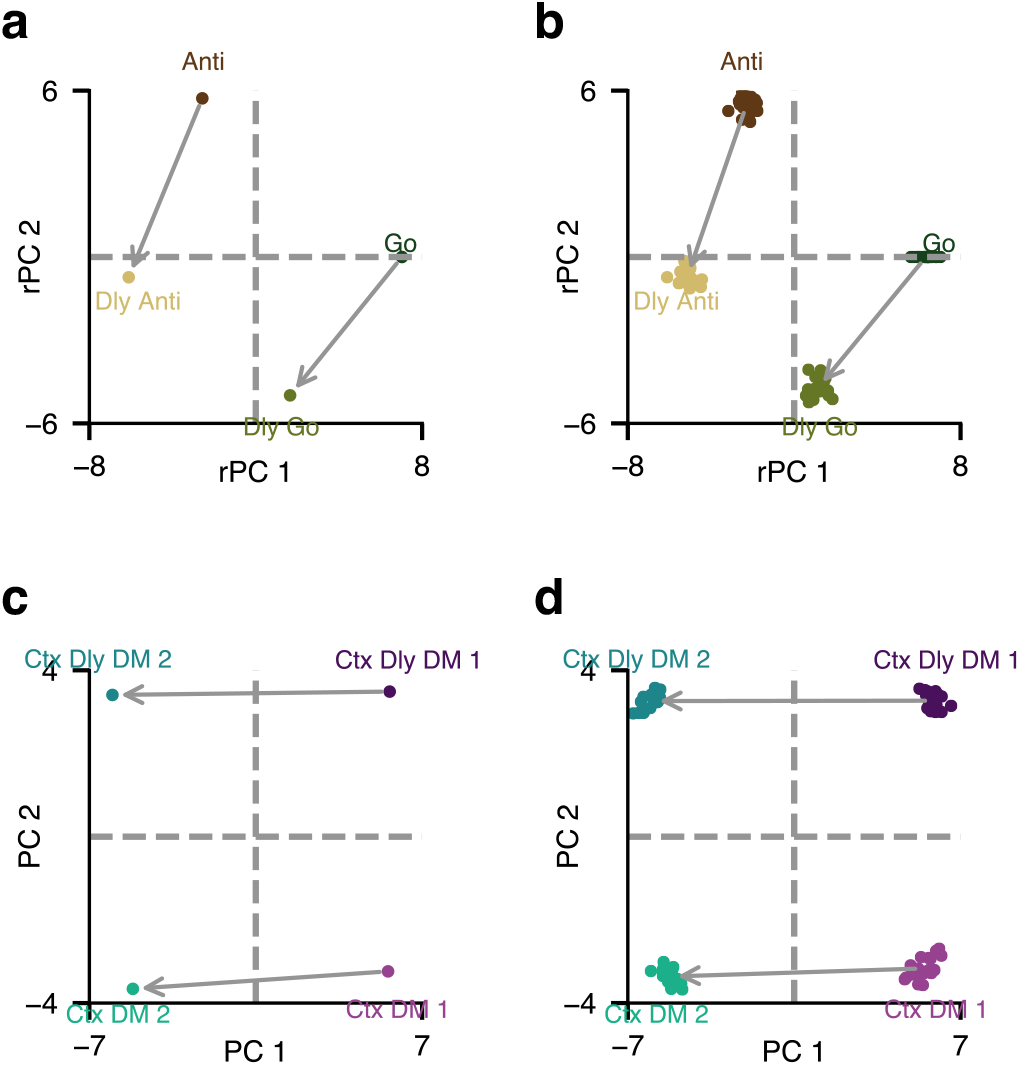
Visualization of connection weights of rule inputs. **(a)** Connection weights from rule input units representing Go, Dly Go, Anti, Dly Anti tasks visualized in the space spanned by the top two principal components (PCs) for a sample network. Similar to 5, the top two PCs are rotated and reflected (rPCs) to form the two axes. **(b)** The same analysis as in **(a)** is performed for 20 networks, and the results are overlaid. **(c)** Connection weights from rule input units representing Ctx DM 1, Ctx DM 2, Ctx Dly DM 1, and Ctx Dly DM 2 tasks visualized in the top two PCs for a sample network. **(d)** The same analysis as in **(c)** for 20 networks.

